# Creeping yeast: a simple, cheap, and robust protocol for the identification of mating type in *Saccharomyces cerevisiae*

**DOI:** 10.1101/2020.01.19.911990

**Authors:** Samantha D. M. Arras, Taylor R. Hibbard, Lucy Mitsugi-McHattie, Matthew A. Woods, Charlotte E. Johnson, Andrew Munkacsi, Sylvie Hermann-Le Denmat, Austen R. D. Ganley

## Abstract

*Saccharomyces cerevisiae* is an exceptional genetic system, with genetic crosses facilitated by its ability to be maintained in haploid and diploid forms. Such crosses are straightforward if the mating type/ploidy of the strains are known. Several techniques can be used to determine mating type (or ploidy), but all have limitations. Here we validate a simple, cheap and robust method to rapidly identify *S. cerevisiae* mating types. When cells of opposite mating type are mixed in liquid media, they “creep” up the culture vessel sides, a phenotype that can be easily detected visually. In contrast, mixtures of the same mating type or with a diploid simply settle out. The phenotype is robust to different media, cell densities, temperatures and strains, and is observable for several days. Microscopy suggests that cell aggregation during mating is responsible for the phenotype. Yeast knockout collection analysis identified 107 genes required for the creeping phenotype, with these being enriched for mating-specific genes. Surprisingly, the RIM101 signalling pathway was strongly represented. We propose that RIM101 signalling regulates aggregation as part of a wider, previously-unrecognized role in mating. The simplicity and robustness of this method makes it ideal for routine verification of *S. cerevisiae* mating type, with future studies required to verify its molecular basis.

## Introduction

The ability to maintain *Saccharomyces cerevisiae* in haploid and diploid forms as well as to perform genetic crosses is fundamental to the success of this yeast as a model organism for molecular genetics (Guthrie & Fink, 1991, Botstein & Fink, 2011). *S. cerevisiae* has two mating types, **a** and α. In nature *S. cerevisiae* is predominately inbreeding, as it routinely switches mating-type to overcome mating-type incompatibility (Haber, 2012). However, in most laboratory *S. cerevisiae* strains mating type is extremely stable and heritable due to deletion of the *HO* endonuclease gene that facilitates this mating type switching (Herskowitz & Jensen, 1991). Mating type is determined by which of two idiomorphs (termed as such because of differences in length and sequence), *MAT***a** or *MAT*α, is present at the *MAT* locus (Haber, 2012). The *MAT*α idiomorph encodes two genes that specify the α-specific phenotype, while the *MAT***a** idiomorph encodes a transcriptional repressor that, in **a**/α diploids, suppresses expression of haploid-specific genes. Haploid cells respond to the mating type pheromones produced by the opposite mating type, triggering a signalling cascade that eventually results in morphological changes that drive cell fusion and karyogamy (Bardwell, 2005).

Mating between compatible *S. cerevisiae* cells in liquid cultures involves an aggregation process that physically holds cells together. This sexual aggregation is mediated by **a** and α agglutinin proteins that are expressed on the surface of the cell wall of haploid **a** and α strains, respectively (Banderas *et al*., 2016). The **a**-agglutinin is composed of two subunits encoded by the *AGA1* and *AGA2* genes, while the α-agglutinin is encoded by *SAG1* (Lipke & Kurjan, 1992). The agglutinin proteins adhere once they come into contact with each other like a sort of molecular Velcro that promotes cell-cell contact (Lipke & Kurjan, 1992). Agglutinins are constitutively expressed at low levels, with their expression being up-regulated in response to mating-type pheromone to facilitate cell aggregation (Lipke & Kurjan, 1992). The sexual aggregation mediated by these agglutinins is likely the first step for mating of yeast cells mixed in suspension (Banderas *et al*., 2016).

Performing genetic crosses in *S. cerevisiae* relies on knowing the mating types of the haploid strains involved. While the mating type and ploidy of established laboratory strains are generally known, there are a number of scenarios where they need to be determined. This includes when dissecting ascospores to produce haploid strains, checking the mating of strains that lack convenient markers, or when mutating the *HO* locus in wild-type strains. Currently, several methodologies are used to determine mating type and ploidy. Flow cytometry can be used to determine if a cell is haploid or diploid, but does not distinguish mating type. A simple method to determine mating type or ploidy is to spread cells following mating on selective medium to look for cells containing markers from both parents, although this obviously requires appropriate markers (Sherman, 1991). A PCR method utilizes *MAT* locus primers that produce different amplicon sizes depending on which mating type idiomorph is present (Bradbury *et al*., 2006). Microscopic observation of the ‘shmoo’ phenotype, which is a manifestation of morphological changes induced upon detection of the opposite mating type pheromone, is also used to identify mating type (Duntze *et al*., 1970, Sprague, 1991, Merlini *et al*., 2013) following mixing of known *MAT***a** or *MAT*α strains with the unknown strain(s). However, shmoo formation can be transient and is only observed in a small number of cells. Finally, the “halo” method, which relies on inhibition of certain strains by pheromone production, can be used to determine mating type but requires tester strains with specific genotypes (Sprague, 1991). Thus, although there is a diversity of methods, they are time-consuming, require specific components, and/or can produce inconsistent or transient results.

Here, we demonstrate a simple, cheap and robust method to enable the rapid identification of the mating type and ploidy of *S. cerevisiae* strains. The principle of the method is that when cells of opposite mating type are mixed, they “creep” up the sides of the culture vessel in an easily observable manner, unlike mixtures involving the same mating type or a diploid strain, which simply settle out. An unknown strain can be mixed with known strains of each mating type in a culture medium, left overnight, and simply observed to allow accurate distinction of both mating types as well as of diploid strains. The response is long-lasting, does not require specialized equipment or reagents, and requires minimal hands-on time. Investigation of the molecular basis for this ‘creeping’ phenotype using microscopy and the yeast knockout collection suggests that it is a consequence of cell aggregation, and indicates a surprising role of the RIM101 pathway in *S. cerevisiae* mating.

## Methods

### Yeast strains and standard growth conditions

All strains used in this study are listed in **Supplementary Table 1**. Culture media used were YPD (1% yeast extract, 2% bacto-peptone and 2% glucose), YPGlycerol (1% yeast extract, 2% bacto-peptone and 2% glycerol), and YNB (0.45% yeast nitrogen base with ammonium sulphate, appropriate amino acids and bases, and 2% glucose (Treco & Lundblad, 1993); **Supplementary Table 2**). Strains were stored in 15% glycerol at −80°C until use.

### Mating type assay

Strains were revived from glycerol stocks on YPD medium then grown overnight in appropriate liquid medium (YPD unless stated otherwise) at 30°C. Strains were diluted to OD_600nm_ 0.2 in the same medium and added in equal volumes to a tube (2 mL per strain for a test tube, 250 μL for a microcentrifuge tube, or 100 μL for a 96-well plate). Cultures were left with no shaking for approximately 18 hrs at ~22°C (room temperature) unless otherwise indicated, after which they were observed and photographed. Time-lapse photography was performed using a Nikon D850 camera with a 60 mm macro lens and polarizing filter. Images were exposed at ISO200, f/16 for 10 s. Photographs were taken every minute from the start of the experiment for 20 hrs. Microscopic observation of cell aggregation was carried out for mating and non-mating cells using a LEICA ICC50 W microscope and the Leica Application Suite (LAS) EZ software (Leica Microsystems v3.2). Observations were performed on 10 μL of 200 μL mating and non-mating cell mixtures from a 96-well plate incubated on the bench with no shaking. Each 10 μL sample was gently collected from the bottom of the well where the creeping phenotype is seen and observed hourly from 2 to 7 hrs following mixing.

### Multiplex PCR to confirm mating type assay

Primers MAT-a 5’-CAATGATTAAAATAGCATAGTCGG-3’, MAT-alpha 5’-CAGCACGGAATATGGGAC-3’ and MAT-R 5’-GGTGCATTTGTCATCCGTC-3’ (Bradbury *et al*., 2006) were used in a multiplex PCR reaction to amplify mating-type specific PCR products from 2 μL of overnight yeast culture (100 μL in YPD) in a final volume of 25 μL. PCR was performed with the KAPA2G Robust DNA polymerase in GC buffer (Custom Science). Cycling parameters were initial incubation of 10 mins at 94°C, then 35 cycles of 94°C for 25 s, 55°C for 25 s and 72°C for 90 s. The *MAT***a** amplicon (489 bp) and *MAT*α amplicon (466 bp) were visualized on a 3% agarose gel run at 100 V for 4 hrs in sodium borate buffer (10 mM NaOH, 36 mM boric acid) and stained with ethidium bromide.

### Genome-wide analysis of the mating type assay

The genetic basis for the “creeping” phenotype was investigated via crosses of BY4742 (*MAT*α) with the gene deletion library (*MAT***a**; BY4741 as parental strain) in round bottom 96-well plates. First, the gene deletion library (Open Biosystems) was incubated overnight at 30°C in 200 μL YPD, and BY4742 was incubated overnight at 30°C in 5 mL YPD. The next day, 80 μL of BY4742 cells (1.26 × 10^7^ cells/mL) was mixed with 120 μL of gene deletion library strains, incubated overnight at 30°C, and imaged using a digital camera. Mating and non-mating phenotypes were identified through independent visual inspection by two people.

### Phenotype enrichment

Genes required for the “creeping” phenotype were analyzed for phenotype enrichment using YeastEnrichr (Chen *et al*., 2013, Kuleshov *et al*., 2016). Functional enrichment for phenotype was evaluated using text-mining associations of each gene with phenotypes reported in publications in PubMed. Significant enrichment was defined for categories with an adjusted *P*-value, a z-score (expected:observed ratio for each phenotype), and a combined score (the product of the z-score and negative logarithm of the *P*-value).

### Spatial analysis of functional enrichment

Genes required for the “creeping” phenotype were annotated for placement in a functionally annotated genetic interaction network (Costanzo *et al*., 2016) via spatial analysis of functional enrichment (SAFE; Baryshnikova, 2016).

### Network analysis

Genes required for the “creeping” phenotype were annotated for function via network- and pathway-based community partitioning as previously described (Busby *et al*., 2019). Using NetworkAnalyst (Zhou *et al*., 2019), genes were mapped in a first-order network within an established yeast global interaction network (the STRING interactome; Szklarczyk *et al*., 2015) with a confidence cutoff of 900 and requirement for experimental evidence. Significant community modules (*P* < 0.05) were identified using the InfoMap algorithm, and pathways enriched in these modules were determined using KEGG pathway (Kanehisa & Goto, 2000, Kanehisa *et al*., 2021) analysis (FDR < 0.05).

## Results and Discussion

### A quick and reliable assay for determination of mating type in *S. cerevisiae*

Routine observations in our laboratory showed that *S. cerevisiae* cells of different mating types settled in non-shaking cultures differently to cells of the same mating type. The manifestation of this mating type phenotype is that cultures of *S. cerevisiae* cells of opposite mating types consistently “creep” up the sides of the culture vessel when mixed and left overnight on the bench with no shaking (**Figure 1**). This “creeping” phenotype is not observed in cultures with a mixture of cells of the same mating type (**a** or α), or where one or both cultures are diploid - these instead simply settle to the bottom of the vessel (**Figure 1**).

**Figure 1:**
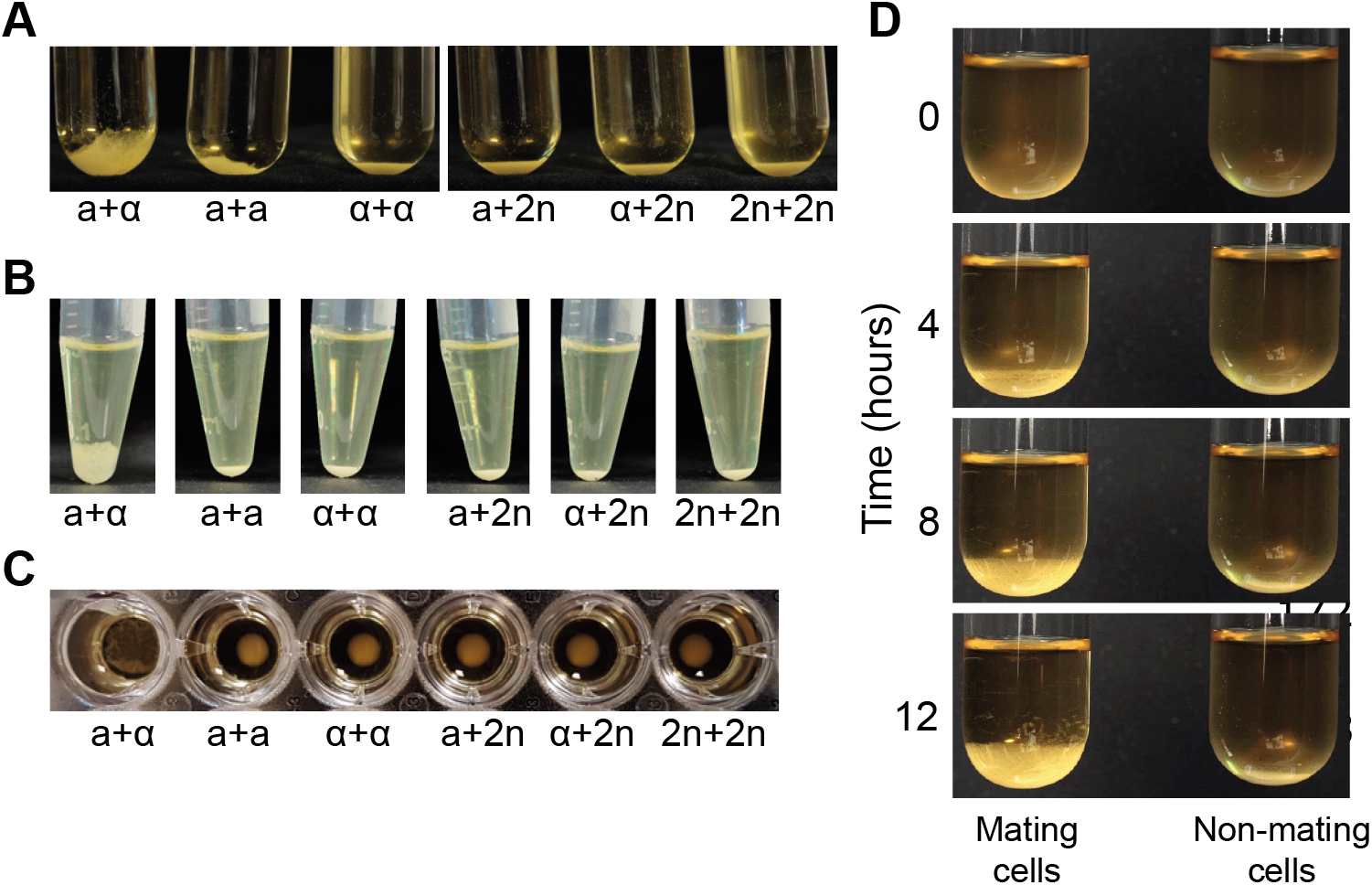
A 50:50 mixture of a+α cells reveals a creeping phonotype. YPD cultures (OD_600nm_ 0.2) were mixed in equal volumes of: 2 mL each in 20 mL glass tubes (**A**); 250 μL each in microcentrifuge tubes (**B**); and 100 μL each in a round-bottomed 96-well plate (**C**). Mixtures were allowed to settle on the bench for 18 hrs and photographed. The creeping phenotype is only observed for the mixtures of opposite mating types (**a**+α). Mating types **a** (a) and α (a), and diploid cells (2n) are indicated below the tubes. The phenotype was not observed in flat-bottomed 96-well plates. **(D)**. Time-lapse photographs of an **a**+α assay at 0, 4, 8, and 12 hrs show the formation of the creeping phenotype compared to a tight pellet for non-mating cells. For full time lapse over 20 hrs, see **Supplementary Figure 1**, and for a video, see **Supplementary Movie**.

We decided to investigate this observation further using a variety of vessels to determine whether it could form a robust mating type assay. We found that the creeping phenotype is observed in glass test tubes, plastic microfuge tubes, and round bottom microwell plates (**Figure 1A-C**), demonstrating its scalability and flexibility, although we did not observe the phenotype in flat-bottomed microwell plates. Time-lapse photography revealed that the formation of the creeping phenotype is visible ~4 hrs after the mixing of YPD cultures of **a** and α cells at room temperature, and intensifies between about 4 and 8 hrs (**Figure 1D**; see **Supplementary Movie** for a complete video of the formation of the creeping phenotype over 20 hrs). Moreover, the phenotype is visible for several days. These results show that this assay is a simple way to distinguish *MAT***a** and *MAT*α strains (formation of the creeping phenotype with one or other of the known mating type tester strains), as well as diploid strains (no creeping phenotype with either known mating type tester strain), and that the assay can be performed in a variety of culture vessel formats.

### Determination of conditions compatible with the mating type assay

We next wanted to characterize how robust the mating type phenotype is under different conditions to assess its practicality as a mating type assay. We first tried different media, including rich and synthetic media, and water. The creeping phenotype forms reliably in both rich and synthetic media containing glucose as the carbon source, but is less obvious when glycerol is used and does not manifest in water (**Figure 2A**). Next we tested the effects of cell density and partner ratio on the phenotype for cells grown in glucose rich medium. Cell density can be varied from 0.2 to 1.8 (OD_600nm_) without affecting the formation of the creeping phenotype (**Figure 2B**), while increasing the proportion of one partner over 60% impedes the creeping phenotype (**Figure 2C**). The assay is symmetric, in that disruption of the creeping phenotype is observed equally regardless of which partner (**a** or α) is present in excess (**Supplementary Figure 2**). Interestingly, detection of the creeping phenotype is improved in 96-well plates compared to tubes (compare **Supplementary Figure 2** to **Figure 2C**). Finally, the creeping phenotype is obvious at temperatures in the optimal growth range for *S. cerevisiae* (~22°C and 30°C), but is nevertheless still visible when the cultures are incubated at 37°C (**Figure 2D**). However, as for cells in water, formation of the creeping phenotype does not occur at 4°C (**Figure 2D**).

**Figure 2:**
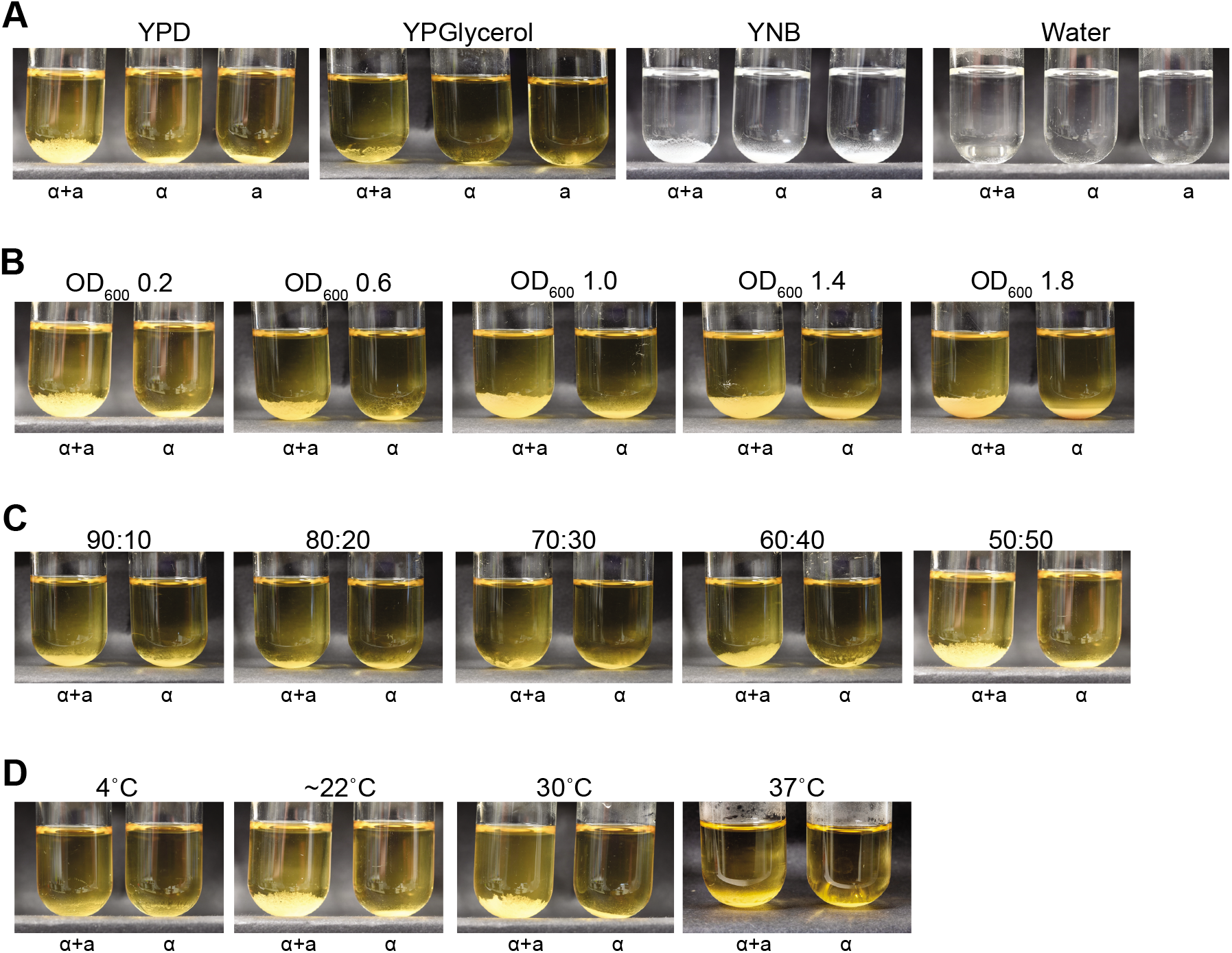
Establishing working parameters for the mating assay. Four parameters were varied: medium (**A**), cell density (**B**), partner ratio (**C**), and temperature (**D**). Strains were diluted to OD_600_ 0.2 in YPD, mixed equally (50:50) and incubated at room temperature (~22°C) unless otherwise indicated. Mating type **a** (a) and mating type α (α) cells are indicated below the tubes. All mixtures were photographed at 18 hrs. (**A**) Overnight YPD cultures were diluted in the indicated media or water prior to the mating assay. Note that YNB and water appear this color because of medium composition, not black and white photography. (**B**) The same overnight YPD cultures from (**A**) were diluted to OD_600nm_ 0.2, 0.6, 1.0, 1.4, or 1.8 to perform the assay. **(C)** Opposite mating type cells were mixed at the different ratios indicated in a final volume of 4 mL (ratios represent α:**a**; see **Supplementary Figure 2** for the reciprocal ratios). (**D**) Mixtures were incubated at 4°C, room temperature, 30°C or 37°C for 18 hrs. Note that the first panel in (**A**) and in (**B**), the last panel in (**C**), and the second panel in (**D**) are all the same.

### Mating type determination of different strains of *S. cerevisiae*

To assess how robust the assay is to strain type, fourteen *S. cerevisiae* strains derived from different wild isolates (Cubillos *et al*., 2009) that include representatives of each of the five major, non-mosaic *S. cerevisiae* clades described by Liti *et al*. (2009) were selected (**Supplementary Table 1**). To test whether the assay is able to determine their mating types, each strain was individually mixed with known tester **a** and α strains using standard conditions (YPD medium, OD_600nm_ 0.2, overnight incubation on the bench without shaking). The length of incubation time necessary to unambiguously visualize the creeping phenotype varied depending on the growth rate of the strain. The mating type determinations made using this assay matched that recorded for each strain, with one exception (**Figure 3A; Supplementary Figure 3**). The one exception is illuminating: we found a discrepancy between the mating type determined by us and that expected (strain 1; *MAT*α by our assay but expected to be *MAT***a**; **Supplementary Table 1**). To check this strain and validate the mating type determinations, we carried out multiplex PCR on seven strains with primers previously designed to determine the presence of the **a** or α idiomorph within the active *MAT* locus (Bradbury *et al*., 2006). Each strain gave the 489 bp **a** or 466 bp α band expected from the creeping phenotype assay (**Figure 3B**), including the discrepant strain. Therefore, this discrepant strain is *MAT*α as the creeping phenotype suggests, with the discrepancy likely a strain designation error rather than a problem with the assay. Together, these results show that the assay can accurately determine mating type for a variety of *S. cerevisiae* strains under a variety of conditions.

**Figure 3:**
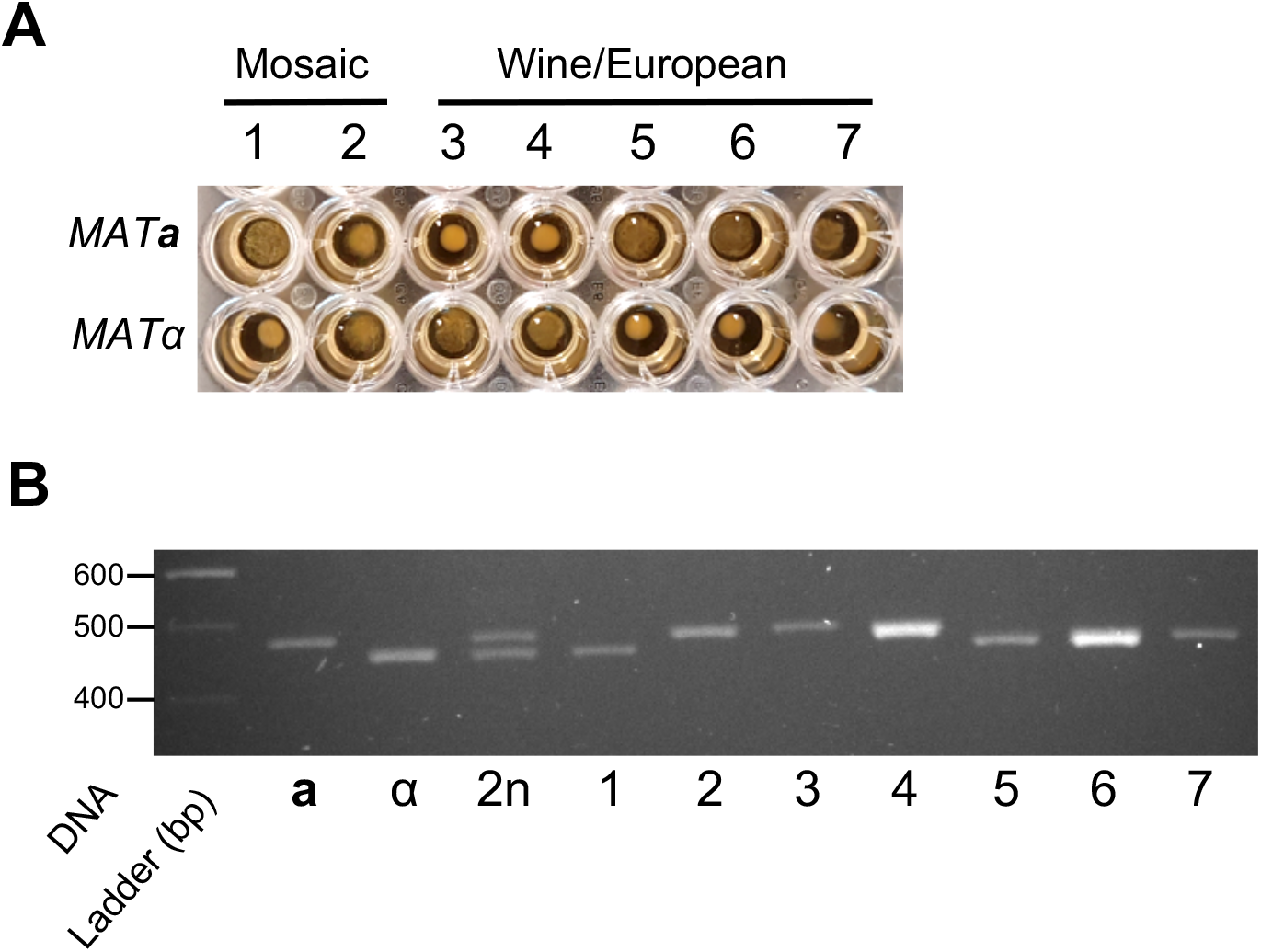
Determination of mating type of wild *S. cerevisiae* strains. **(A)** Seven strains from the Mosaic and Wine/European clades identified in Liti *et al*. (2009) were mixed with *MAT***a** and *MAT*α tester strains using standard assay conditions (YPD medium, OD_600nm_ 0.2, room temperature incubation). Mixtures were photographed after 18 hrs. The creeping phenotype is visible for strains 1, 5, 6 and 7 crossed with *MAT***a** tester but not with *MAT*α tester. The opposite result is visible for strains 2, 3 and 4. (**B**) Mating assay results were confirmed by multiplex PCR carried out on cells from the seven strains and tester **a**, α and 2n strains. Expected *MAT***a** and *MAT*α specific amplicons are 489 bp and 466 bp, respectively. As expected, both amplicons are observed in the diploid strain. Seven additional strains are shown in **Supplementary Figure 3**.

### Sexual aggregation is implicated in the creeping phenotype

To shed light on the basis of the creeping phenotype, we made microscopic observations of the cells as the cultures began to creep. These observations show increased aggregation of cells with time when both mating types are present, but not when only one mating type is present (**Figure 4; Supplementary Figure 4**). Therefore, these results implicate aggregation as the basis for the creeping phenotype. This is consistent with the nature of the phenotype (cells “moving” up the vessel sides through adherence and growth), the requirement for a medium supporting growth, the poor performance on glycerol (which is not a favored carbon source for *S. cerevisiae*), the absence of the phenotype at 4°C (sexual aggregation has previously been reported to be cold sensitive (Zhao *et al*., 2001)), and the phenotype becoming evident from ~4 hrs (around the time of mating initiation (Duntze *et al*., 1970, Sena *et al*., 1973)).

**Figure 4:**
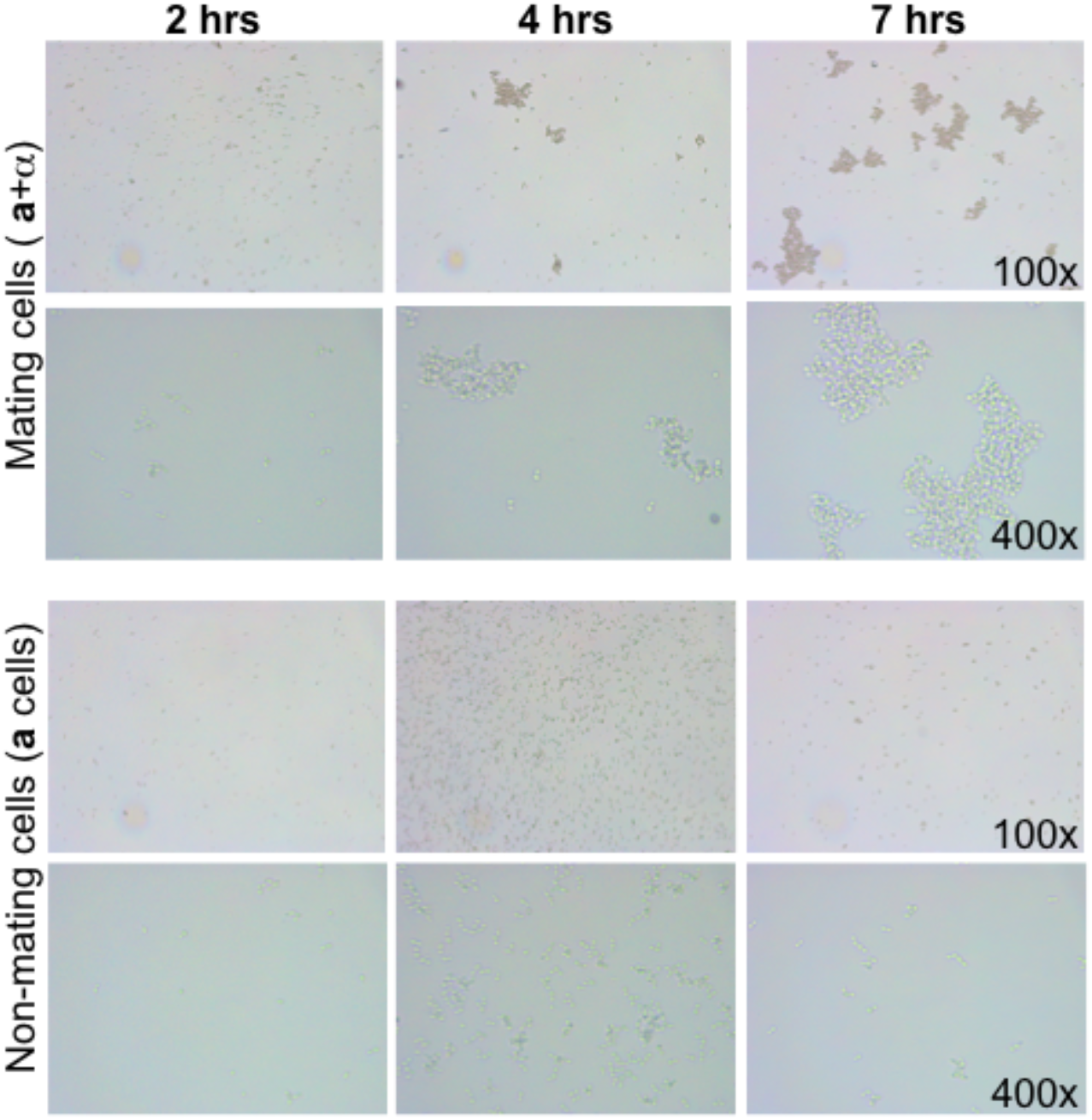
Creeping cells aggregate over time. Mating and non-mating samples of **a**+α cells or haploid **a** cells were observed under the microscope. Images were taken from 2 to 7 hrs after cells were mixed. Representative images at 100x and 400x magnification are shown. Extensive cell aggregation is observed for mating cells from 4 hrs following mixing, while no appreciable aggregation is observed for non-mating cells over the same time period. See **Supplementary Figure 4** for the full time course.

### Functional genomics confirms the creeping phenotype is associated with mating

To gain further insight into the mechanism behind the creeping phenotype, we used a yeast deletion library (Giaever & Nislow, 2014) to examine the impact of non-essential gene deletions on the ability to produce the creeping phenotype. To do this, we mated 4,294 haploid gene deletion strains in a *MAT***a** background void of any growth defect with a wild-type *MAT*α strain, performing two biological replicate experiments. From this we identified 107 gene deletion strains that failed to produce the creeping phenotype in either replicate (**Supplementary Table 3**). These 107 genes comprised 84 characterized genes, and 23 uncharacterized open reading frames (ORFs) that encode putative proteins, dubious ORFs, or proteins of unknown function.

As expected, the 107 genes required for the creeping phenotype are strongly enriched for genes involved in yeast mating. This conclusion is supported by the results of an unbiased assessment of the genetics underlying the creeping phenotype. This was performed by using YeastEnrichr (Kuleshov *et al*., 2016) to identify over-represented phenotypes associated with the 107 genes compared to random gene sets. The top four functions (by adjusted *P*-value < 0.05) are all involved in mating (mating efficiency, mating response, pheromone production and pheromone sensitivity), as are other highly enriched functions such as nuclear fusion during mating and shmoo formation (**Supplementary Table 4**). Moreover, most of the genes reported to be involved in mating signalling (Merlini *et al*., 2013) were amongst our 107 gene set, although none of the genes involved in cell-cell fusion were (**Table 1**). This enrichment in mating signalling but not cell fusion is consistent with the creeping phenotype resulting from aggregation, which occurs before fusion. Surprisingly given the deletion collection is *MAT***a**, *STE2* that encodes the receptor for pheromone α (Burkholder & Hartwell, 1985) was not amongst our 107 genes (**Table 1**). However, it has been reported that strains deleted for *STE2* display a low background of mating (Jahng *et al*., 1988), which might explain the persistence of the creeping phenotype in absence of *STE2*. Conversely, the inclusion of *MFA1* amongst our 107 genes (**Table 1**) is surprising considering that *MFA1* and *MFA2* both encode the **a** pheromone and were shown to be functionally redundant (Michaelis & Herskowitz, 1988). This might suggest that *MFA2* is not able to compensate for the absence of *MFA1* whereas *MFA1* can compensate for the absence of *MFA2* to produce the creeping phenotype, which is consistent with *MFA2* producing three-fold less **a** pheromone than *MFA1* (Chen *et al*., 1997). Finally, consistent with the evidence implicating aggregation, we found that the creeping phenotype was not detectable in the absence of the *AGA1* or *AGA2* agglutinin genes (**Supplementary Table 3**). Thus, as expected, the creeping phenotype seems to depend on yeast mating genetic pathways and the genetic results are consistent with the involvement of mating aggregation.

**Table 1.** Overlap between mating signalling/cell-cell fusion genes and genes required for the creeping phenotype

| Function | Gene | Note |
| --- | --- | --- |
| **SIGNALLING COMPONENTS** |  |  |
| Pheromone **a** | *MFA1<sup>a</sup>, MFA2\** |  |
| Pheromone $\alpha$ | *MF(ALPHA)1<sup>b</sup>*<br>*MF(ALPHA)2* | $\alpha$ specific gene<sup>c</sup><br>$\alpha$ specific gene |
| G-protein coupled receptors |  |  |
| Receptor for pheromone **a** | *STE3* | $\alpha$ specific gene |
| Receptor for pheromone $\alpha$ | *STE2\** | |
| G-protein $\alpha$ subunit | *GPA1* | |
| G-protein $\beta$ subunit | *STE4* | |
| G-protein $\gamma$ subunit | *STE18<sup>d</sup>* | Not in the YKO library |
| PAK kinase | *STE20* |  |
| MAPK scaffold | *STE5* |  |
| MAPK scaffold | *STE50* |  |
| MAPKKK | *STE11* |  |
| MAPKK | *STE7* |  |
| MAPK | *FUS3, KSS1* |  |
| Transcription factor | *STE12* | Not in the YKO library |
| Scaffold for shmoo orientation | *FAR1* |  |
| Cdc42 GTPase | *CDC42* | Essential gene |
| Cdc42-GEF | *CDC24* | Essential gene |
| Cdc42, Ste5 & Ste20-scaffold | *BEM1* |  |
| Ras GTPase | *RAS1* |  |
| Formins | *BNI1, BNR1* |  |
| **FUSION COMPONENTS** |  |  |
| Transmembrane Proteins | *PRM1, FIG1, KEX2* |  |
| Cell wall remodelling | *FUS1, FUS2* |  |
| Lipid raft protein | *RVS161* |  |
| Type V myosin | *MYO2* | Not in the YKO library |
| Tropomyosin | *TPM1* |  |

<sup>a</sup> genes required for the creeping phenotype are shown in red
<sup>b</sup> genes whose deletion does not affect the creeping phenotype are shown in black <sup>c</sup> $\alpha$ specific genes are not expected to affect the creeping phenotype in the *MATa* gene deletion YKO library context
<sup>d</sup> genes that were not present (either missing or essential) in the deletion library are shown in grey
\* genes whose deletion unexpectedly does not affect the creeping phenotype; see text for discussion

### Functional enrichment and network analyses implicate RIM101 signalling in the creeping phenotype

To shed more light on the genetic basis for the creeping phenotype, we performed two further enrichment analyses on our set of 107 genes. First, we used spatial analysis of functional enrichment (SAFE) to investigate how genetic interactions involving the 107 genes relate to the 19 interaction domains previously identified in the yeast genetic interaction landscape (Baryshnikova, 2016, Costanzo *et al*., 2016). This revealed a concentration of genetic interactions in six functional domains: multivesicular body (MVB) sorting/RIM signalling, mitochondria, chromatin, transcription, metabolism, and cytokinesis (**Figure 5A**).

**Figure 5:**
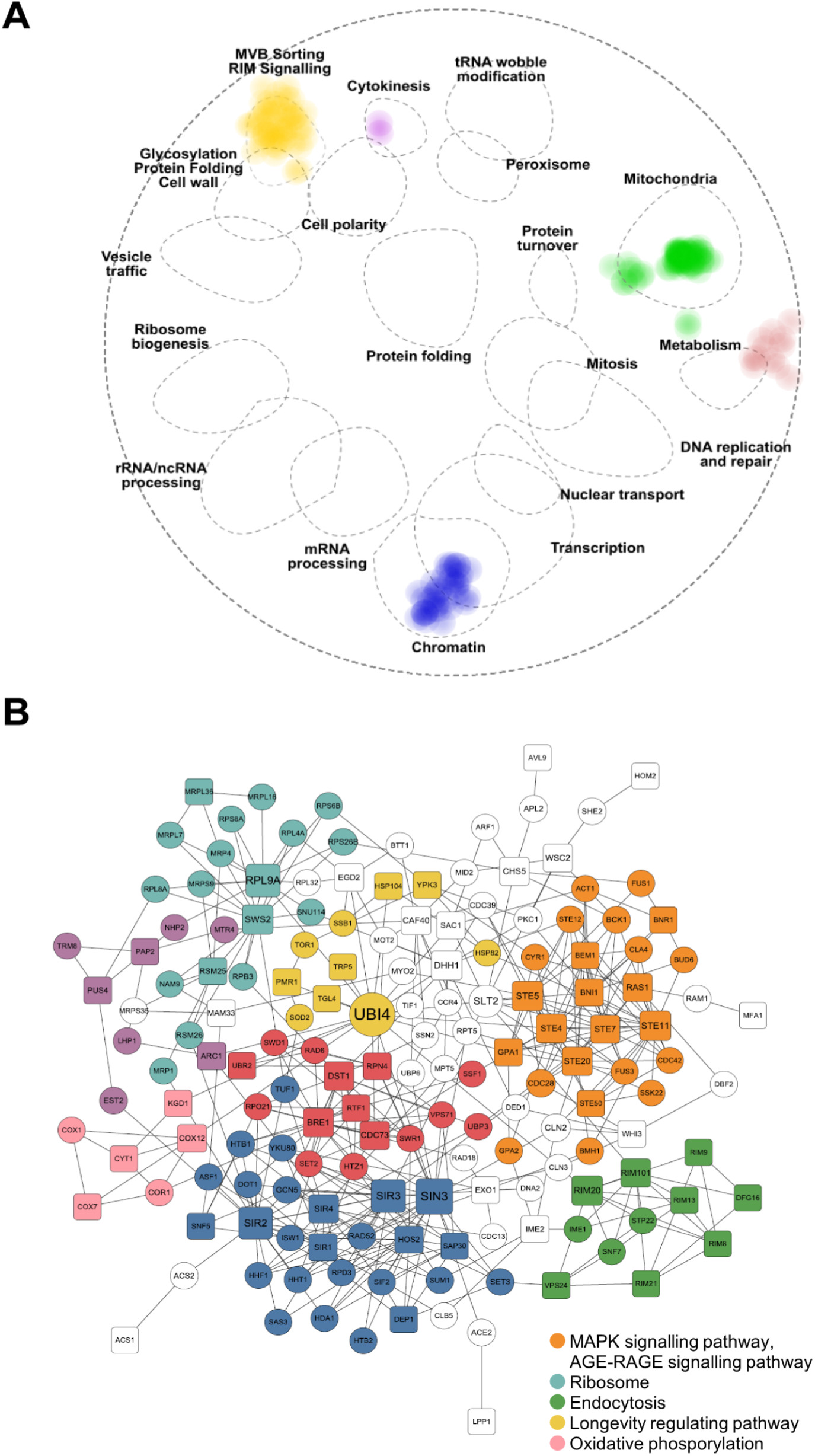
Enrichment analyses of genes required for the creeping phenotype. **(A)** Spatial analysis of functional enrichment (SAFE) of the 107 deletion strains required for the creeping phenotype. All 107 genes were mapped in an established genetic interaction network to identify over-represented regions. The 107 genes show significant membership to six of the previously defined 19 functional regions (Baryshnikova, 2016, Costanzo *et al*., 2016), illustrated by dotted ellipses, with the intensity of color proportional to the number of genes. (**B**) Pathway enrichment for first-order minimum network analysis of the genes required for the creeping phenotype. 65 of the 107 genes appear in this network. Genes required for creeping phenotype are shown as square nodes; interacting genes as circular nodes. The first-order network was generated in NetworkAnalyst (Zhou *et al*., 2019) with the STRING interactome database where nodes and edges represent genes and genetic/physical interactions, respectively. Node size is based on betweenness score, which is indicative of node importance for overall network connectivity. Nodes are colored by module analysis representing tightly clustered subnetworks with more internal connections than randomly expected in the whole network. Nodes not assigned to a module are without color. The most significantly enriched pathway (lowest *P*-value less than 0.05) in the KEGG database was identified for each module. Modules indicated in the legend at bottom right had significant pathway enrichment (*P* < 0.05), while those not listed in the legend did not.

Second, we used NetworkAnalyst (Zhou *et al*., 2019) to map the 107 genes in a first-order network within an established global network comprised of interactions from the STRING interactome (Szklarczyk *et al*., 2015). This classified the 107 genes into an interactive network connecting 65 of the 107 genes with eight statistically significant modules (*P* < 0.05) representative of subnetworks that are more interactive than expected at random (**Figure 5B**). To evaluate over-representation of pathways in these modules, an enrichment analysis of interactions for association to the KEGG pathway database was conducted. Five pathways were significantly enriched for five of the modules (**Figure 5B**): MAPK signalling (*P* = 1.9 × 10^−20^), ribosome (*P* = 5.4 × 10^−7^), endocytosis (*P* = 5.0 × 10^−4^), longevity regulating (*P* = 7.3 × 10^−3^), and oxidative phosphorylation (*P* = 1.6 × 10^−5^). These analyses also show enrichment of mating pathways. For example, the MAPK signalling module (**Figure 5B**) contains many hallmark mating genes (*e.g*., *STE4, STE5, STE7, STE11, STE20, STE50, GPA1, BNI1, BNR1, BEM1*, and *RAS1*), and is integral to the mating pathway (Bardwell, 2005; **Table 1**). Thus, these modules can provide functional insight into the creeping phenotype.

The strongest and most surprising enrichment in both enrichment analyses was for RIM101 signalling (auto-annotated in NetworkAnalyst as “endocytosis”, presumably because the RIM101 pathway contains ESCRT proteins; **Figure 5**). This enrichment is striking: of the twelve genes in the RIM101 pathway, nine are amongst our 107 gene set (**Figure 6**). Eight of these appear in the NetworkAnalyst analysis (**Figure 5B**). The exception is *YGR122W*. which is required for the creeping phenotype but fell below the NetworkAnalyst protein-protein interaction confidence threshold. *IME1* does not show as required for the creeping phenotype, but, although appearing in NetworkAnalyst endocytosis module, is not considered part of the core RIM101 pathway (Lorenz & Cohen, 2014; **Figure 6**). The remaining two genes, *SNF7* and *SPT22*, were not present in the gene deletion library. Thus, all the core RIM101 pathway genes we could assess are required for the creeping phenotype.

**Figure 6:**
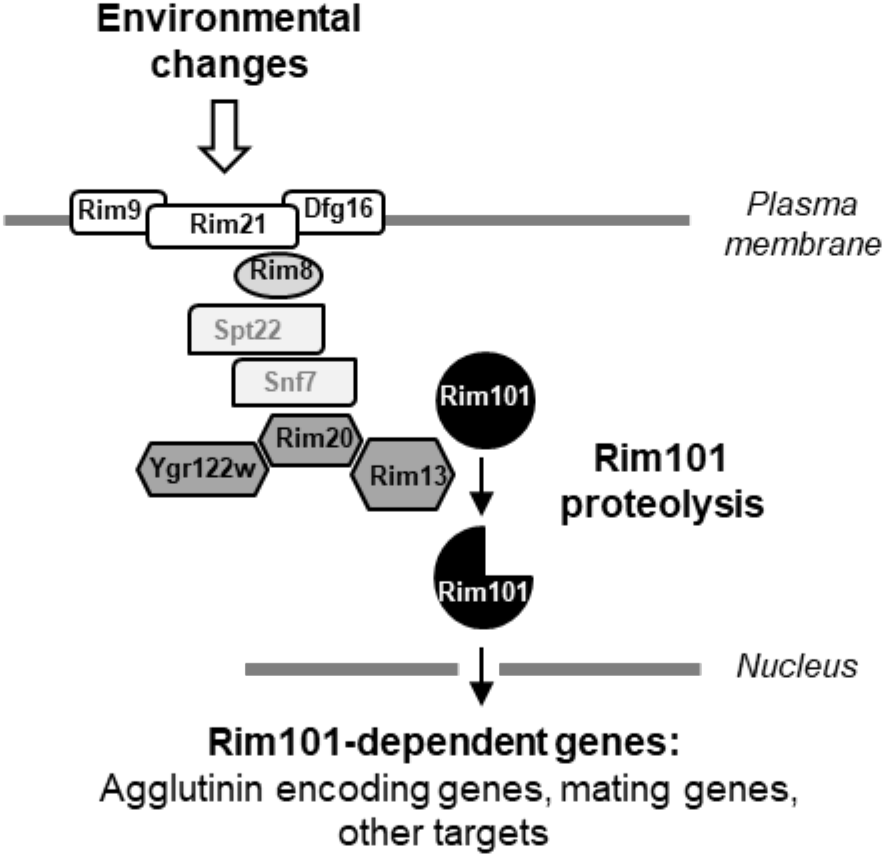
Schematic representation of the RIM101 signalling pathway and its potential involvement in *S. cerevisiae* mating. The fungal-specific RIM101 signalling pathway is based on the proteolytic processing of the Rim101 transcription factor. The integral membrane proteins Rim21, Dfg6 and Rim9 constitute a surface response complex that recruits the arrestin-like Rim8 at the plasma membrane. Downstream ESCRT (<u>E</u>ndosomal <u>S</u>orting <u>C</u>omplex <u>R</u>equired for <u>T</u>ransport) proteins Spt22, Vsp24, Snf7 are recruited in a Rim8-dependent manner to constitute a platform on which the proteolytic complex (Ygr122w, Rim20, Rim13) forms. The calpain-like cysteine protease Rim13 then cleaves Rim101, inducing its nuclear localization and cellular transcriptional response. We propose that Rim101 targets include genes responsible for the sexual agglutination response amongst other genes that mediate mating. Apart from *SNF7* and *SPT22*, which were not present in our gene deletion library screen, all genes in the RIM101 pathway were found to be required for the creeping phenotype (**Figure 5B**, **Supplementary Table 3**). Adapted from Ost *et al*. (2015).

The RIM101 pathway is a conserved fungal-specific signalling pathway that mediates adaptation to external pH (Penalva & Arst, 2004). It is named after the pH-responsive transcription factor Rim101 whose activation by C-terminal proteolytic cleavage is the key output of the pathway (Orejas *et al*., 1995, Li & Mitchell, 1997; **Figure 6**). Besides its role in environmental pH response, the pathway is involved in diverse cellular behaviors that include sporulation and haploid invasive growth in *S. cerevisiae* (Su & Mitchell, 1993, Li & Mitchell, 1997), salt tolerance in *S. cerevisiae* and *Candida albicans* (Lamb & Mitchell, 2003), pathogenicity in *C. albicans, Aspergillus fumigatus* and *Cryptococcus neoformans* (Davis, 2009), and cell wall assembly in *S. cerevisiae* and *C. neoformans* (Castrejon *et al*., 2006, O’Meara *et al*., 2013, Zhao *et al*., 2019).

Involvement of the RIM101 pathway in the creeping phenotype of *S. cerevisiae* mating cells strongly suggests that this signalling pathway also plays an important role in the first step(s) of mating, particularly in mating partner aggregation (**Figure 4**, **Supplementary Figure 4**). We hypothesize that the RIM101 pathway mediates this process by regulating expression of **a** and α agglutinin encoding genes, given Rim101 is a transcription factor. While we are not aware of the RIM101 pathway having previously been proposed to mediate mating in *S. cerevisiae, rim* mutants have been reported to display mating defects (Castrejon *et al*., 2006). Direct support for our hypothesis comes from results showing that deletion of *RIM101* and *RIM13* (encoding the protease responsible for Rim101 activation; **Figure 6**) both result in downregulated expression of *AGA2* and *MFA1* (which encode the **a**-agglutinin adhesive subunit and pheromone **a**, respectively) in a *MAT***a** strain (Lamb & Mitchell, 2003; **Supplementary Table 5**). In addition, when we examined a publicly-available genome-wide RNA-seq dataset where *RIM101* is deleted in a *MAT*α strain (Read *et al*., 2016), we found that the α-agglutinin and pheromone a encoding genes (*SAG1*, *MF(ALPHA)1, MF(ALPHA)2*) were significantly downregulated, mirroring the impact of *RIM101* deletion in a *MAT***a** strain (**Supplementary Table 5**). These data also show downregulation of *AGA1*, encoding the **a**-agglutinin anchor subunit in the *MAT*α context (**Supplementary Table 5**), which is likely an outcome of *AGA1* being expressed in both *MAT***a** and *MAT*α cells (Lipke & Kurjan, 1992). Thus, Rim101 seems to regulate expression of mating aggregation genes in both mating types. Moreover, expression of *RIM101* is absolutely required in both partners for mating in the yeast *Yarrowia lipolytica* (Lambert *et al*., 1997), although the mating was assessed on solid rather than liquid medium. Finally, **Supplementary Table 5** shows that around 2/3rds (19 out of 30) of the genes reported to be involved in mating signalling (Merlini *et al*., 2013) have their expression significantly affected by *RIM101* deletion, suggesting a wider involvement of the RIM101 pathway in mating. Altogether, our data and existing evidence suggest a novel and direct role for the *RIM101* signalling pathway in sexual agglutinin expression as part of a more general role in mating signalling, thereby explaining the involvement of this pathway in the creeping phenotype.

## Conclusions

Here we describe a rapid, inexpensive and robust method to determine not only the mating type of an unknown *S. cerevisiae* strain, but also whether the strain is haploid or diploid. While we serendipitously discovered the creeping phenotype in our laboratory, we have heard anecdotally that we are not the first to discover it and that it is currently used as a mating type assay in some laboratories. Thus, here we focused on evaluating and optimizing the phenotype to provide it as an alternative mating type assay for the yeast community. The assay is as simple as mixing an “unknown” sample with known *MAT***a** and *MAT*α strains at roughly equal cell densities and observing the results after 18 hours, thus it is cheap, robust, and requires minimal hands-on time. So far, all *S. cerevisiae* strains we have tried in our laboratory are amenable to the assay, but we do not know whether it will work for all *S. cerevisiae* strains or for other *Saccharomyces* species.

Our microscopy data suggest the phenotype results from cell aggregation that occurs during yeast mating. This conclusion is supported by the timing of the creeping phenotype, which occurs around the start of mating following mixing of compatible strains and is concomitant with visible signs of aggregation. It is also consistent with the requirement for conditions facilitating growth, and in particular with the longevity of the phenotype - well past when mating is likely to have completed. Moreover, demonstration that the creeping phenotype depends on many characterized mating genes, and in particular on the *AGA1* and *AGA2* agglutinin genes, is also consistent with aggregation underlying the phenotype.

Surprisingly, our functional genomics screen identified the RIM101 pathway as being necessary for the creeping phenotype. All components of this pathway we could assess (all three surface response complex genes *RIM9, RIM21*, and *DFG16*; two of the four of Rim8-dependent platform component genes (*RIM8, VSP24*); and all thee proteolytic complex genes *RIM13, RIM20, YGR122w*) are required for the creeping phenotype. The gene encoding the transcription factor Rim101 is also required, proteolytic activation of which is the key output of the pathway. We suggest that Rim101 regulates the agglutinin genes, thus dependence of the creeping phenotype on the RIM101 pathway can be explained simply by a decrease of the expression of these genes in the absence of Rim101 activation. Our examination of published data supports this contention, and a more general, previously-unrecognized role for RIM101 signalling in *S. cerevisiae* mating. While Rim101-dependent regulation of the agglutinin genes can explain the dependence of the creeping phenotype on the RIM101 pathway, the trigger that leads to Rim101 activation during mating is unknown. Thus, future work will be required to confirm whether Rim101 directly regulates the agglutinin genes and other mating genes, and how the RIM101 pathway is activated during mating. The simplicity of assaying the creeping phenotype will aid these future approaches, as well as further investigations into the phenotype.

## Acknowledgements

We thank Ganley lab members for comments on the manuscript.

