## Supplementary Figure 1 for "Creeping yeast: a simple, cheap, and robust protocol for the identification of mating type in *Saccharomyces cerevisiae*"

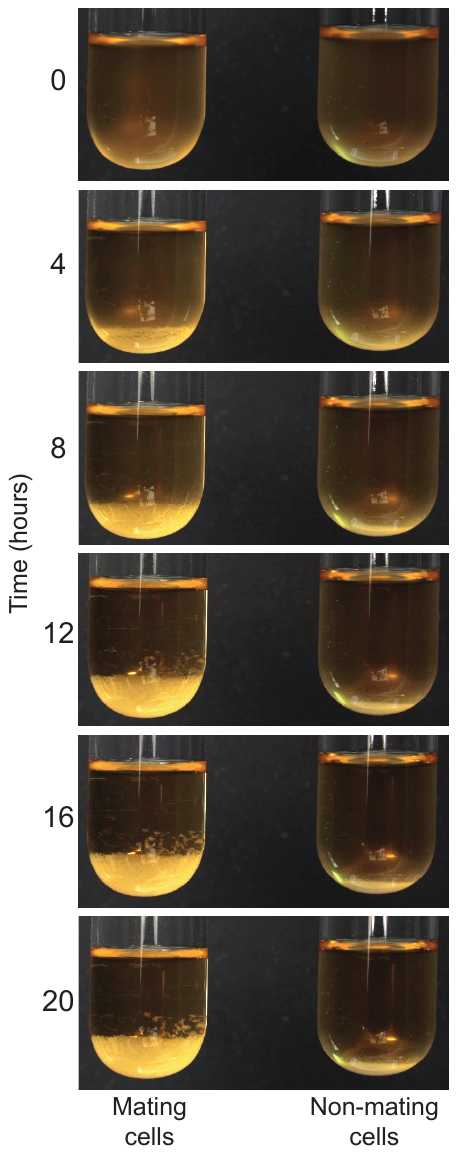

**Supplementary Figure 1: Time-lapse photography of the mating assay.** An **a**+α assay was set up as described in **Figure 1A** and photographs were taken every minute for 20 hours. Views at 0, 4, 8, 12, 16 and 20 hours show the formation and persistence of the creeping phenotype compared to a tight pellet for non-mating cells. For a video of the full time-lapse, see **Supplementary Movie.**
