## Supplementary Figure 2 for "Creeping yeast: a simple, cheap, and robust protocol for the identification of mating type in *Saccharomyces cerevisiae*"

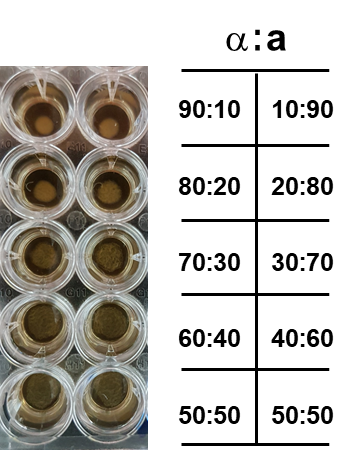

**Supplementary Figure 2: Symmetric formation of creeping phenotype with different ratios of each mating type.** Opposite mating type cells were mixed at the different ratios indicated in a final volume of 200 μL in a 96 well plate. The plate was photographed after 18 hours at 22˚C. The right column shows increased ratios in favour of mating type **a** cells, the left column shows increased ratios in favour of mating type α cells. This assay supplements **Figure 2C.**
