## Supplementary Figure 3 for "Creeping yeast: a simple, cheap, and robust protocol for the identification of mating type in *Saccharomyces cerevisiae*"

North American

*MAT****a***

*MATα*

**8	9	10	11	12	13	14**

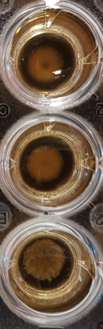

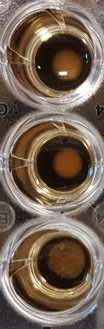

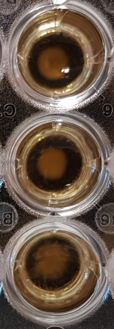

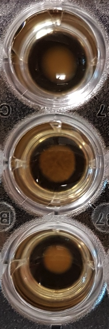

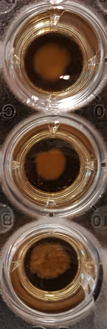

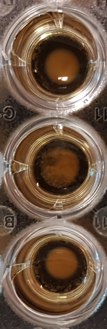

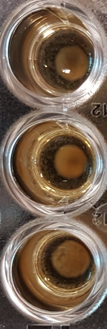

West African

Malaysian

Sake

Strain

**Supplementary Figure 3:** Mating assay as in **Figure 3A** for wild isolates that include four additional representatives of major, non-mosaic *S. cerevisiae* clades described by Liti *et al.,* 2009. Assays photographed after 5 days of incubation at 22°C. The bottom wells (Strain) contain strains 8 to 14 only. All strains are described in **Supplementary Table 1.**
