## Supplementary Figure 4 for "Creeping yeast: a simple, cheap, and robust protocol for the identification of mating type in *Saccharomyces cerevisiae*"

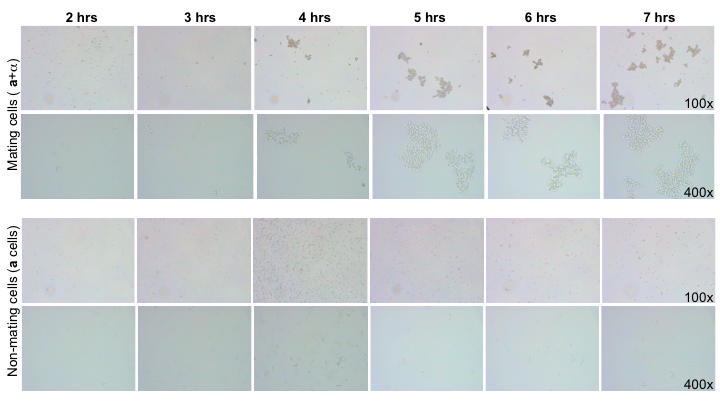

**Supplementary Figure 4: Creeping cells aggregate over time.** Full time course of mating and non-mating samples of **a**+α cells or haploid **a** cells observed under the microscope as described in **Figure 4**. Images were taken from 2 to 7 hrs after cells were mixed. Representative images at 100x and 400x magnification are shown.
