## Supplementary Table 1 for "Creeping yeast: a simple, cheap, and robust protocol for the identification of mating type in *Saccharomyces cerevisiae*"

**Supplementary Table 1:** Strains used in this study

| **Strain** | **Name** | **Genotype** | **Clade^a^** | **Reference** |
| --- | --- | --- | --- | --- |
| haploid *MAT***a** | NOY408-1b | *MAT****a*** *ade2-1 ura3-1 his3-11 trp1-1*  *leu2-3, 112 can1-100* | Mosaic | Nogi et al. 1991 |
| haploid *MAT*α | NOY408-1a | *MAT*α *ade2-1 ura3-1 his3-11 trp1-1*  *leu2-3, 112 can1-100* | Mosaic |  |
| diploid (2n) | NOY398 | *MAT****a****/ MAT*α *ade2-1/ ade2-1 ura3-1/ ura3-1 his3-11/ his3-11 trp1-1/ trp1-1 leu2-3, 112/ leu2-3, 112 can1-100/can1-100* | Mosaic |  |
| 1^b^ | 3582/ UWOPS87-2421 | *MAT***a**, ∆*ho::HygMX*, ∆*ura3::KanMX* | Mosaic | Cubillos et al. 2009 |
| 2 | 3589/ UWOPS83-787.3 | *MAT***a**, ∆*ho::HygMX*, ∆*ura3::KanMX* | Mosaic |  |
| 3 | 3593/ YJM981 | *MAT***a**, ∆*ho::HygMX*, ∆*ura3::KanMX* | Wine/European |  |
| 4 | 3597/ DBVPG6765 | *MAT***a**, ∆*ho::HygMX*, ∆*ura3::KanMX* | Wine/European |  |
| 5 | 3616/ BC187 | *MAT*α*,* ∆*ho::HygMX*, ∆*ura3::KanMX* | Wine/European |  |
| 6 | 3620/ DBVPG1373 | *MAT*α, ∆*ho::HygMX*, ∆*ura3::KanMX* | Wine/European |  |
| 7 | 3624/ L-1528 | *MAT*α, ∆*ho::HygMX*, ∆*ura3::KanMX* | Wine/European |  |
| 8 | 3625/ DBVPG6044 | *MAT*α, ∆*ho::HygMX*, ∆*ura3::KanMX* | West African |  |
| 9 | 3626/ NCYC110 | *MAT*α, ∆*ho::HygMX*, ∆*ura3::KanMX* | West African |  |
| 10 | 3627/ UWOPS03-461.4 | *MAT*α, ∆*ho::HygMX*, ∆*ura3::KanMX* | Malaysian |  |
| 11 | 3603/ UWOPS05-217.3 | *MAT***a**, ∆*ho::HygMX*, ∆*ura3::KanMX* | Malaysian |  |
| 12 | 3630/ Y12 | *MAT*α, ∆*ho::HygMX*, ∆*ura3::KanMX* | Sake |  |
| 13 | 3607/ YPS128 | *MAT***a**, ∆*ho::HygMX*, ∆*ura3::KanMX* | North American |  |
| 14 | 3632/ YPS128 | *MAT*α, ∆*ho::HygMX*, ∆*ura3::KanMX* | North American |  |
| haploid *MAT*α | BY4742 | *MAT*α *his3Δ1 leu2Δ0 met15Δ0 ura3Δ0* | Derivative of S288c lineage | Open Biosystems |
| haploid *MAT***a** | BY4741  YKO parental strain^c^ | *MAT***a** *his3Δ1 leu2Δ0 met15Δ0 ura3Δ0* | Derivative of S288c lineage | Open Biosystems |

^a^ According to Liti et al, 2009.

^b^ Strain 3582/ UWOPS87-2421 is reported as being mating type **a**, but we have shown that our version of this strain is mating type α, possibly as a result of a mis-labelling error.

^c^ Yeast Knockout (YKO) library used in this study.
